# Measuring Transcription Factor Binding and Gene Expression using Barcoded Self-Reporting Transposon Calling Cards and Transcriptomes

**DOI:** 10.1101/2021.04.15.439516

**Authors:** Matthew Lalli, Allen Yen, Urvashi Thopte, Fengping Dong, Arnav Moudgil, Xuhua Chen, Jeffrey Milbrandt, Joseph D. Dougherty, Robi D. Mitra

## Abstract

Calling cards technology using self-reporting transposons enables the identification of DNA-protein interactions through RNA sequencing. Although immensely powerful, current implementations of calling cards in bulk experiments on populations of cells are technically cumbersome and require many replicates to identify independent insertions into the same genomic locus. Here, we have drastically reduced the cost and labor requirements of calling card experiments in bulk populations of cells by introducing a DNA barcode into the calling card itself. An additional barcode incorporated during reverse transcription enables simultaneous transcriptome measurement in a facile and affordable protocol. We demonstrate that barcoded self-reporting transposons recover *in vitro* binding sites for four basic helix-loop-helix transcription factors with important roles in cell fate specification: ASCL1, MYOD1, NEUROD2, and NGN1. Further, simultaneous calling cards and transcriptional profiling during transcription factor overexpression identified both binding sites and gene expression changes for two of these factors. Lastly, we demonstrated barcoded calling cards can record binding *in vivo* in the mouse brain. In sum, RNA-based identification of transcription factor binding sites and gene expression through barcoded self-reporting transposon calling cards and transcriptomes is an efficient and powerful method to infer gene regulatory networks in a population of cells.

## Introduction

Calling cards is a uniquely powerful method to genetically record interactions between a protein of interest and the genome^1,2^. Briefly, a protein of interest is fused to a transposase which can insert a transposon ‘calling card’ into the genome at sites of DNA-protein interaction such as transcription factor binding sites (TFBS). Early protocols recovered inserted transposons from genomic DNA^1^, but a recent technical innovation termed the ‘self-reporting transposon’ (SRT) allows for the facile recovery of calling cards through RNA sequencing (RNA-seq)^3^. RNA-mediated mapping of transposon insertions is more efficient than previous DNA-based protocols, and this protocol enables the simultaneous identification of TFBS and changes in gene expression in single cells^4^. However, in bulk experiments on populations of cells, the RNA-mediated protocol is technically cumbersome, requiring a large number of replicates to identify independent insertions into the same genomic locus^4^. Here, we present two crucial modifications of the SRT technology and protocol to facilitate its use and to enable joint recording of TFBS and gene expression in populations of cells: barcoded SRTs and barcoded transcriptomes.

Current implementations of the mammalian calling card protocol employ a hyper-active *piggyBac* transposase^5^. An inherent constraint of this transposase is its requirement for a ‘TTAA’ tetranucleotide sequence for transposon insertion. As a result, multiple independent calling card insertions often occur at the same genomic location in different cells. Since the identification of TF binding sites is based on transposition count rather than read density, if these independent insertions are not distinguished, it limits the dynamic range of bulk calling card experiments. In the DNA-based calling card protocol, we solved this problem by including a barcode between the terminal repeats of the transposon that could be recovered by inverse PCR. However, this location is not compatible with the more efficient RNA-based protocol due to frequent “barcode swapping” during cDNA amplification^6,7^. As a result, current best practices for calling card experiments require a large number of biological replicates (typically 8-12) for each condition to increase the number of insertions that can be detected at a given TTAA location^4^. While this improves the quantitative readout of these experiments, experimental cost and labor scale linearly with the number of replicates. Therefore, as an alternative approach, we sought to embed a unique barcode within the terminal repeat (TR) of the self-reporting transposon, the best location to enable reliable recovery without barcode swapping. Doing so is challenging, however, because all published sequences of the *piggyBac* transposon TRs are completely invariant, indicating strong sequence constraints on TR function which might preclude barcode insertion^8–12^.

Here, we performed targeted mutagenesis of the *piggyBac* terminal repeat sequence to identify sites that could accommodate barcodes in calling card experiments. We discovered at least four consecutive nucleotides within the TR that were tolerant of a range of mutations without major reductions in transposition efficiency. As a resource to the scientific community, we have developed a set of barcoded *piggyBac* SRT plasmids and modified the calling card analysis software to utilize these barcodes. We demonstrate that barcoded SRT calling cards can map the genomic binding sites of transcription factors (TFs) involved in cell fate specification and transdifferentiation *in vitro*. Additionally, we combined barcoded SRT calling cards with bulk RNA barcoding and sequencing (BRB-seq)^13^. This enables us to simultaneously identify TFBS and accompanying transcriptional changes from multiple TFs in an easy and affordable protocol.

Lastly, we demonstrate that barcoded SRTs facilitate *in vivo* calling card experiments in the mouse brain, reducing labor by 10-fold. These innovations simplify bulk SRT calling card experiments, enable barcoding of experimental conditions, and allow for pooled library preparations that substantially reduce cost and labor. This simple protocol for simultaneously measuring transcription factor binding and gene expression changes will facilitate the inference of gene regulatory networks for TFs involved in development, cellular reprogramming, and disease.

## Results

### Identifying Candidate Regions for Barcode Insertion in piggyBac Terminal Repeat

The SRT consists of a promoter driving a reporter (e.g., fluorescent protein or puromycin resistance cassette) flanked by the transposon terminal repeat sequences (TR), the part of a transposon that is recognized by its cognate transposase. Importantly, there is no polyadenylation (poly(A)) signal sequence after the reporter gene, so gene transcription proceeds through the TR and into the genome. This design allows the SRT to report its genomic location in cellular RNA (Figure 1A)^3^. To maximize compatibility with calling cards library preparation and minimize template switching, the ideal barcode location would be as close to the genomic insertion site as possible. Because sequences outside of the TRs are not inserted into the genome, barcodes cannot be introduced there (Fig. 1A, site 1). A barcode inserted between the reporter gene and the TR, as implemented in our DNA-based calling cards protocol^1^, would be ~300 bp away from the informative transposon-genome junction (Fig. 1A, site 2). This would retain a long stretch of shared sequence present in all amplicons that would lead to extensive barcode swapping during the SRT amplification PCR step in library preparation^6,7^.

**Figure 1:**
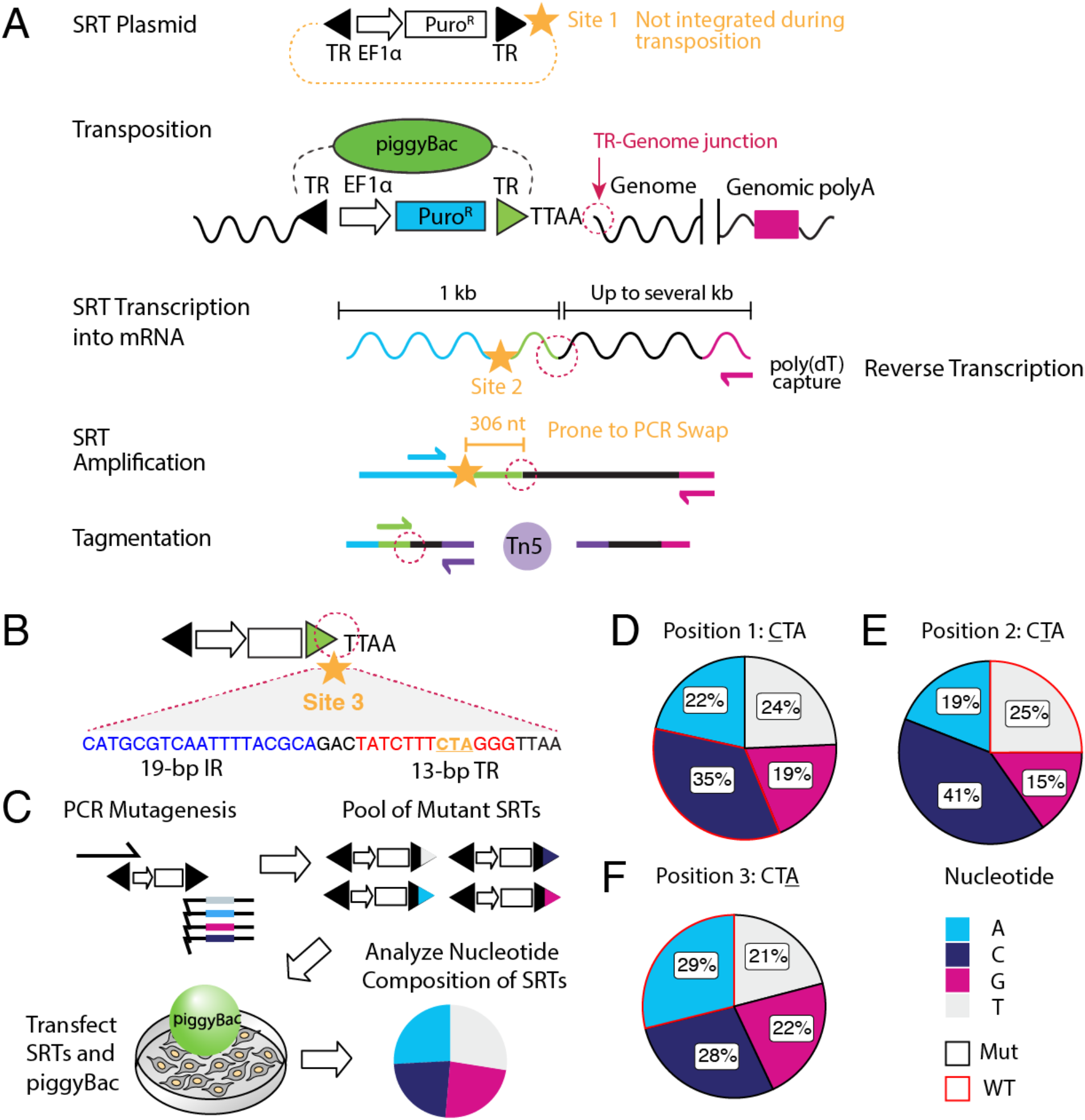
Barcoding the self-reporting transposon. A) Schematic overview of the SRT construct, Calling Card method, and sequencing library preparation. Candidate sites for barcode insertions are indicated with gold stars. The TR-Genome junction, used to map transposon insertions, is circled in dotted magenta line. B) Barcode site 3 is within the *piggyBac* TR sequence, immediately adjacent to the TR-Genome junction. Underlined nucleotides in the 13-bp terminal inverted repeat region (‘CTA’, gold) were targeted for mutagenesis by mutagenic PCR. C) Overview of calling card rapid mutagenesis scheme. Mutant amplicons were transfected into cells with *piggyBac* transposase and integrated calling cards were collected. Nucleotide frequency for each mutagenized position of integrated SRTs were calculated. Nucleotide frequency at D) position 1, E) position 2 and F) position 3 of integrated mutated SRTs. Wild-type sequences are outlined in red. All four possible nucleotides were well-represented at all three mutated positions. IR: internal repeat. TR: terminal repeat. EF1a: eukaryotic translation elongation factor 1 α promoter. SRT: self-reporting transposon. nt: nucleotide. kb: kilobase. Puro^R^: puromycin resistance cassette. WT: wild-type. Mut: mutant.

Therefore, we sought to introduce a barcode into the TR itself (Fig. 1B, site 3), directly adjacent to the TR-genome junction. Such a strategy has two major advantages compared to other approaches. First, a barcode in this position could be captured in the same sequencing read as the transposon-genome junction, simplifying the protocol. Second, by eliminating as much constant intervening sequence as possible, there is little risk of introducing aberrant chimeric PCR products during sequencing library preparation^6,7^. Whereas modifications to TRs from other transposases such as *SleepingBeauty* have been successfully engineered^9^, similar efforts have revealed extensive sequence constraints on *piggyBac* TRs for efficient transposition^10,11^. Nevertheless, we sought to identify candidate regions within the TR that might accommodate a DNA barcode.

The minimal *piggyBac* TR consists of a 19-bp internal repeat (IR), a 3-bp spacer, and a 13-bp terminal invert repeat^12^ (Fig. 1B). These sequences are critical for *piggyBac* recognition, cleavage, and transposition. Notably, all published sequences of the 13-bp terminal invert repeat in the *piggyBac* TR are completely invariant. DNase I footprinting of *piggyBac* binding to its TRs revealed strong binding across much of this region^8^, yet a few bases were less protected and therefore might be a candidate region for inserting a barcode (Fig. 1B, underlined nucleotides, gold).

### Targeted Mutagenesis Generates Mutant SRTs with High Transposition Efficiency

We developed a simple and rapid screening protocol to generate and identify mutant *piggyBac* TR sequences capable of successful transposition (Fig. 1C). We designed primer sequences to introduce single point mutations into our candidate region using PCR. Purified PCR products encoding puromycin-resistance SRTs flanked by mutated TRs were directly transfected into HEK293T cells along with unfused hyper-active *piggyBac*. If mutated amplicons are compatible with transposition, they will be inserted into the genome and confer puromycin resistance. We selected for transposition events after 4 days by adding puromycin. We extracted RNA 3 days after selection, and prepared bulk SRT libraries according to established protocols with modifications described^4^ (Methods).

We sequenced calling card libraries using RNA-seq and mapped genomic transposition events from at least two independently generated mutant SRT pools for each position. Each library yielded 75,000-150,000 unique insertion sites providing a representative view of genomic insertion efficiency for mutant SRTs. Analysis of transposition events revealed that all 3 candidate positions within the *piggyBac* TR accommodated mutations without greatly diminishing transposition ability (Fig. 1D-F). Each of the three mutagenized positions tolerated all 4 nucleotides at similar frequencies, hence generating at least 12 unique transposon barcodes.

Having obtained successful transposition of SRTs with single mutations, we next tested whether multi-nucleotide mutations within this region could be tolerated. Using PCR, we introduced 3 consecutive mixed bases (Ns, where N can be A,C,G, or T) into this region to generate a total of 64 barcoded SRTs. We transfected pools of these mutant SRT PCR amplicons into cells and again prepared calling card libraries after puromycin selection. Analysis of hundreds of thousands of transposition events showed that all 64 mutant transposons could be integrated into the genome, albeit at varying degrees of efficiency (Figure 2A). To better understand sequence preferences governing transposition efficiency, we generated a sequence motif from the top 30 most abundantly inserted transposons. Cytosine was slightly favored in the first two positions, and thymine was strongly disfavored from the third position (Supplementary Figure 1). Among mapped transposition events, we also observed the presence of mutations at a fourth nucleotide position immediately adjacent to our targeted bases, leading us to test whether this position could also be modified. Following the same approach, we generated SRTs with mutations in this position and prepared calling card libraries from two independently transfected sets of cells. As with the other single nucleotide SRT mutants, we found that this position could also tolerate all 4 nucleotides (Fig. 2B).

**Figure 2:**
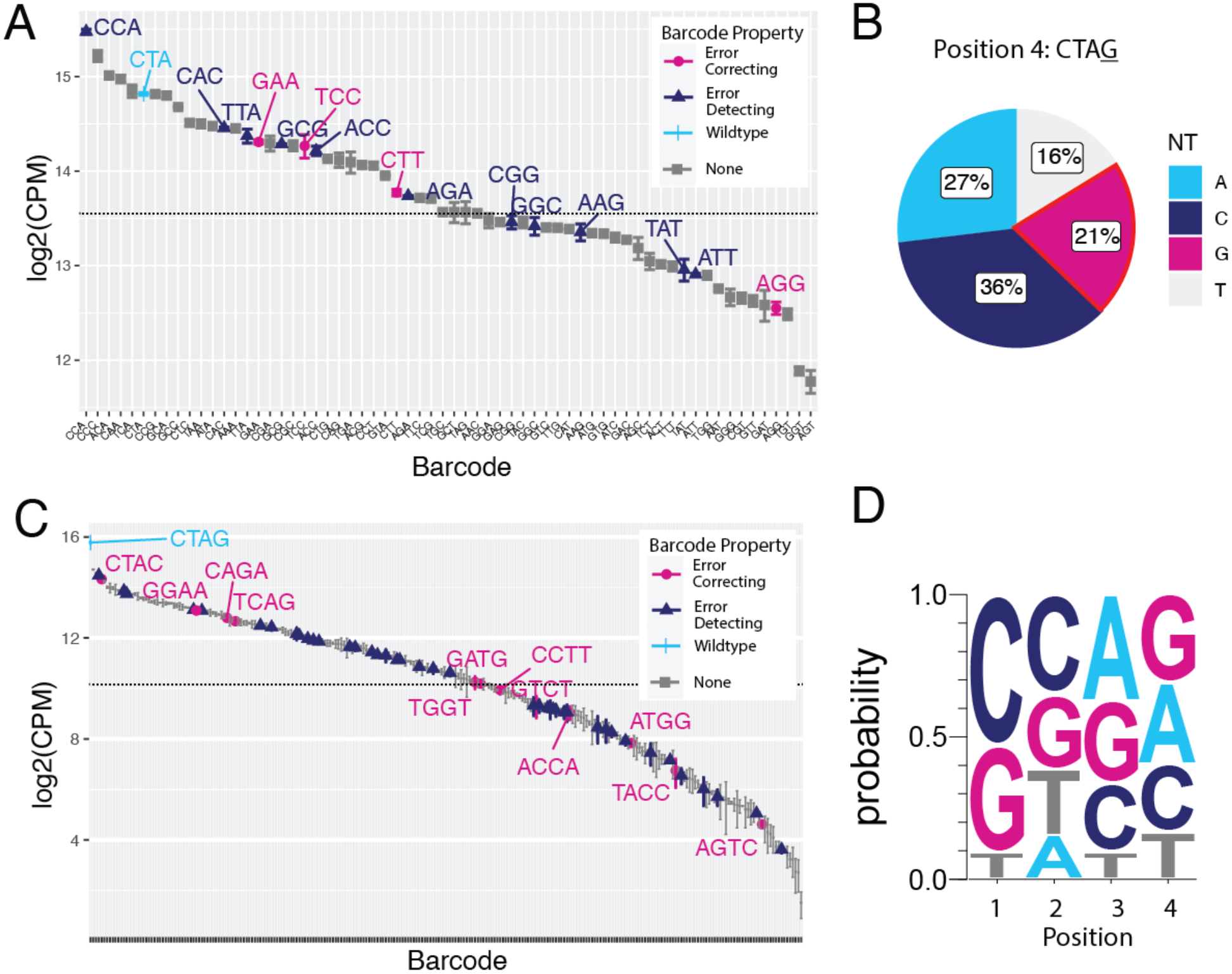
Multi-nucleotide mutagenesis in *piggyBac* terminal repeat discovers integration-competent barcoded SRTs. A) Normalized counts of integration of events for 64 possible combinations of 3 nucleotide barcodes at the targeted region are shown (log2 counts per million (CPM)). All 64 barcoded SRTs could integrate into the genome. Black dotted lines indicate 50^th^ percentile of read counts. Data are plotted as mean and SEM from two independent replicates. B) Targeted mutagenesis at a fourth position in the terminal repeat identified another site that could tolerate all 4 nucleotide substitutions while retaining integration-competence. Wild-type sequence (‘G’) is outlined in red. C) Normalized counts (log2 CPM) of insertions for 256 combinations of 4-nt barcodes. All 256 barcodes were present at varying degrees of insertional efficiency. Wild-type sequence is colored cerulean. Error-correcting and error-detecting barcodes are colored respectively in magenta and midnight blue. D) Sequence logo of the top 100 most abundantly inserted 4-nt barcoded SRTs reveals modest sequence preference for integration efficiency. CPM: counts per million sequencing reads.

Longer barcodes are preferable in sequencing applications as they not only increase the number of unique sequences available but can also have advantageous properties including error detection and error correction^14^. A four-nucleotide barcode yields 256 unique sequences including 40 error-detecting and 12 error-correcting barcodes. To generate a pool of 256 mutant transposons, we introduced 4 consecutive mixed bases (Ns) into the TR using a degenerate primer. We collected and analyzed over 160,000 unique transposition sites in the genome and found all 256 possible mutated transposons were inserted into the genome (Fig. 2C). We analyzed the nucleotide composition of the top 100 most abundantly inserted transposons to reveal sequences mediating transposition efficiency (Fig. 2D). Overall preferences were modest except for a strong favoring of C/G in the first position and a disfavoring of thymine in the third position. These results suggest that a fixed sequence for these 4 nucleotides is not required for binding and transposition by the *piggyBac* transposase.

Given the compatibility of mutations in this region of the TR with transposition, we tested whether we could insert a single nucleotide in this region to further increase barcode length. We generated mutant SRTs with a single nucleotide insertion and performed calling cards with these. We observed that very few cells survived selection and consequently few transposition events were recovered from this experiment. Among the recovered transposition events, the most prevalent sequence matched the wild-type SRT with no insertion. Of the recovered SRTs that did contain an inserted nucleotide, many of the sequences also contained a nearby 1-nt deletion which suggests a strict TR length constraint for successful transposition (Supplementary Table 1). The inserted nucleotide may have disrupted any step of *piggyBac* recognition, cleavage, and transposition by changing the sequence, shape, or flexibility of the transposon^8,15^. Thus, focusing on just the 4 nucleotide barcodes, as a resource to the community, we individually cloned the top 24 integration-competent error-detecting barcodes into two self-reporting vectors. These SRT vectors include an adeno-associated viral (AAV) vector carrying a tdTomato reporter SRT compatible with *in vivo* calling card experiments^16^ and a non-AAV SRT vector encoding the puromycin resistance gene.

### Using barcoded SRTs to map binding sites of transcription factors involved in cell fate specification

To demonstrate that barcoded SRTs facilitate TFBS recording in cellular populations, we performed calling card experiments for four TFs using this method. We chose to record the binding of four members of the basic helix-loop-helix (bHLH) family: Achaete-scute homolog 1 (ASCL1), Myogenic Differentiation 1 (MYOD1), Neuronal Differentiation 2 (NEUROD2), and Neurogenin 1 (here referred to as NGN1). These TFs are implicated in cell fate specification and cellular reprogramming^17–22^. Interestingly, all four TFs recognize the same canonical E-box motif *in vivo*, bind some overlapping and unique sites in the genome, and regulate distinct gene expression programs^23^. To perform calling card experiments, we first created mammalian expression vectors containing fusion proteins of each of the four TFs to the N-terminus of hyperactive *piggyBac* separated by an L3 linker^24^. We transfected HEK293T cells expressing fused or unfused *piggyBac* with wild-type or barcoded versions of SRTs encoding either tdTomato or puromycin-resistance reporters and harvested RNA after ~1 week. We prepared and sequenced SRT calling card RNA-seq libraries and analyzed the data to identify transposon insertions in the genome. Calling card peaks were called as described^1,25^ and analyzed for enriched motifs and neighboring genes using HOMER^26^. We then performed Gene Ontology enrichment analysis on sets of genes located near TFBS^27^.

For each of the four bHLH factors, we recovered hundreds of thousands of genomic insertion events and called thousands of calling card peaks (Supplementary Table 2). Motif enrichment analysis for each factor recovered several enriched bHLH E-box motifs, including the known motifs for Ascl1, MyoD, and NeuroD1 (Figure 3A). This motif recovery suggests barcoded calling cards identified bona fide TFBS for these factors. For NEUROD2, the top 3 enriched motifs belonged to specific neuronal bHLHs including NeuroD itself (Fig. 3A). Likewise for MYOD1, the top 3 enriched motifs belonged to myogenic bHLHs of the MyoD family (Fig. 3A), indicating specificity of the calling card peaks for the TFs of interest. This result supports the interpretation that while the core E-box motif is common to all factors, nucleotides flanking this motif may confer binding specificity^28^. For ASCL1, in addition to recovering bHLH motifs, we also observed an enrichment of Jun/Fos and other basic zipper (bZIP) motifs which might indicate the binding of additional TFs at these sites.

**Figure 3:**
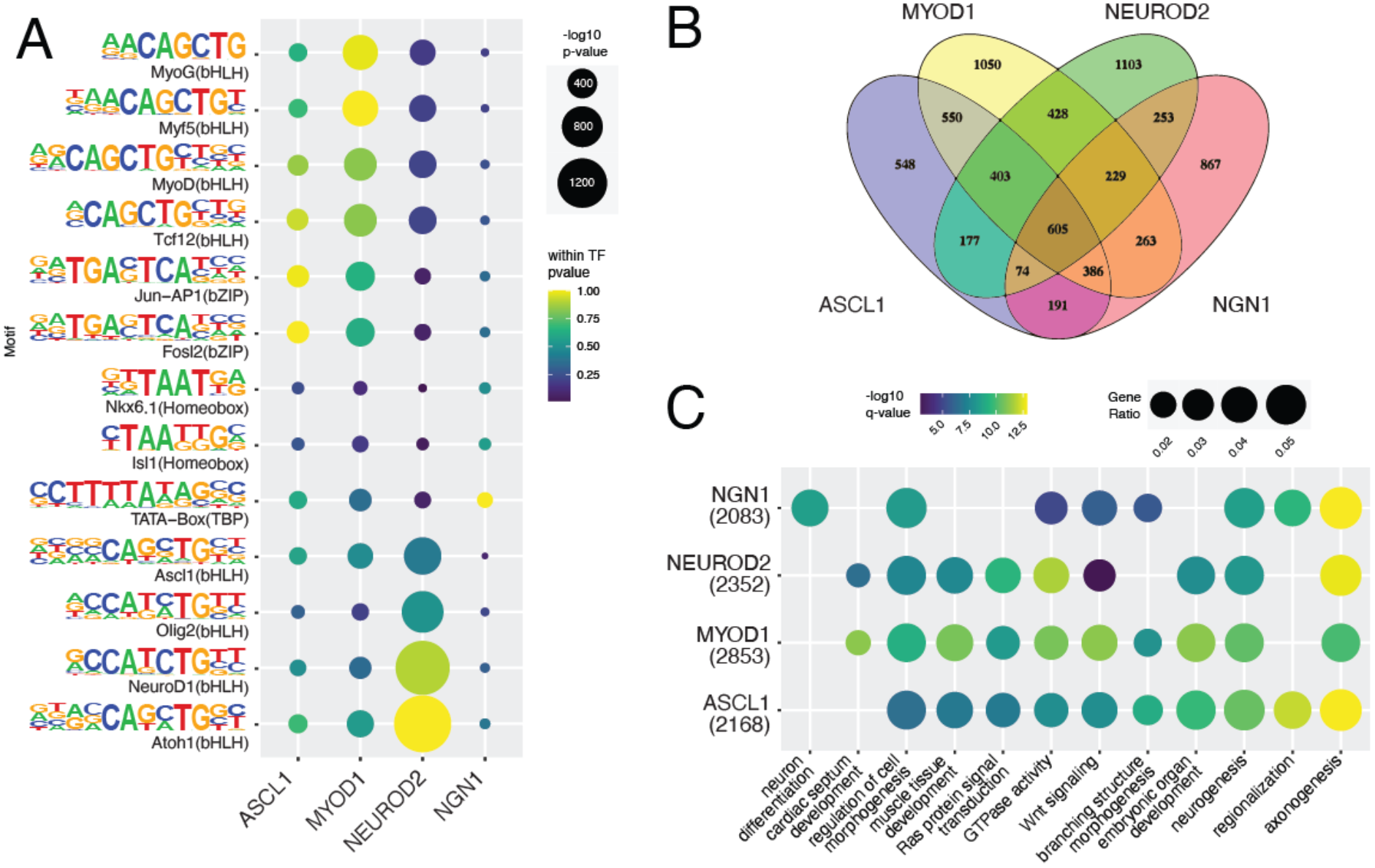
Calling cards using barcoded SRTs recover known binding motifs for bHLH factors near genes related to known TF functions. A) Top binding motifs for each motif were retrieved from DNA sequences in calling card peaks. These sites are enriched for the canonical E-box motif as well as bHLH TFs including or related to each TF. B) Venn diagram of genes proximal to called peaks for each TF indicates both shared and distinct binding of these TFs. C) Gene Ontology enrichment analysis reveals terms related to neurogenesis and myogenesis. bHLH: basic helix-loop-helix. bZIP: basic zipper.

Next, we identified genes located near each TFBS, and characterized the shared and differential binding of bHLH TFs ^29^. Consistent with their recognition of a common E-box motif, all four TFs bound near many of the same genes (Fig. 3B, Supplementary Fig. 2A-B). While many genes were overlapping across TFs, each TF also had its own set of unique genes near TFBS. To gain insight into the regulatory roles of these TFs, we performed Gene Ontology enrichment analysis on sets of genes located near TFBS identified by barcoded SRTs^27^. Gene Ontology terms identified for genes proximal to the neurogenic TFs ASCL1, NEUROD2, and NGN1 were enriched for neuronal pathways including axonogenesis and neuron projection development, reflecting their known roles in neuronal reprogramming (Fig. 3C)^20,30,31^. MYOD1 binding sites were located near genes strongly enriched for roles in cardiogenesis and muscle development (Fig. 3C, Supp. Fig. 2C). Consistent with prior findings of MYOD1 binding some neuronal targets^18^, we found some enrichment for binding at genes enriched for neurogenic pathways. The observed enrichment of neuronal and muscle genes is particularly notable given the calling card assay was performed in human embryonic kidney cells which do not natively express any of the assayed TFs. That all factors are able to recognize and bind specific genes enriched for their known functions implies either a permissive binding environment in HEK293T cells or cell-type independent target access by these TFs. This also highlights that subtle differences in nucleotide sequences flanking the common core E-box motif can confer binding specificity at functionally distinct gene sets^28^.

### Barcoded SRTs and Transcriptomes Enable Simultaneous Mapping of TFBS and Gene Expression

SRTs were specifically invented to enable simultaneous readout of gene expression and transcription factor binding in single cells^3^, but can also be used to map TFBS in populations of cells as demonstrated here and previously^3,16^. Because SRTs are amplified from poly(A) RNA, we sought to prepare SRT and poly(A) mRNA sequencing libraries in parallel from the same sample. To reduce cost and labor of library preparation, multiple poly(A) mRNA-seq libraries can be barcoded during reverse-transcription then pooled for library preparation and sequencing^13^. We have previously modified this barcoding protocol to employ the 10x Genomics single cell 3’ v2 chemistry which enables turnkey analysis of RNA-seq data using CellRanger^32,33^.

To facilitate simultaneous preparation of SRT calling card and poly(A) 3’ RNA-seq libraries, we introduced a sample barcode and unique molecular identifier (UMIs) into the poly(dT) capture oligonucleotide (Figure 4A). Each experimental replicate is reverse transcribed using an oligo(dT) capture oligo with a unique sample barcode, then multiple samples can be pooled for ultra-affordable transcriptomic analysis^13^. Calling card experiments could also be designed such that experimental replicates use distinctly barcoded SRTs (individual barcodes or sets of barcodes) so that the same pool of cDNA can then be used to amplify SRTs and mRNA in parallel reactions. Otherwise, SRT libraries can be amplified individually. Sequencing libraries of amplified products are then prepared by tagmentation^34^.

**Figure 4:**
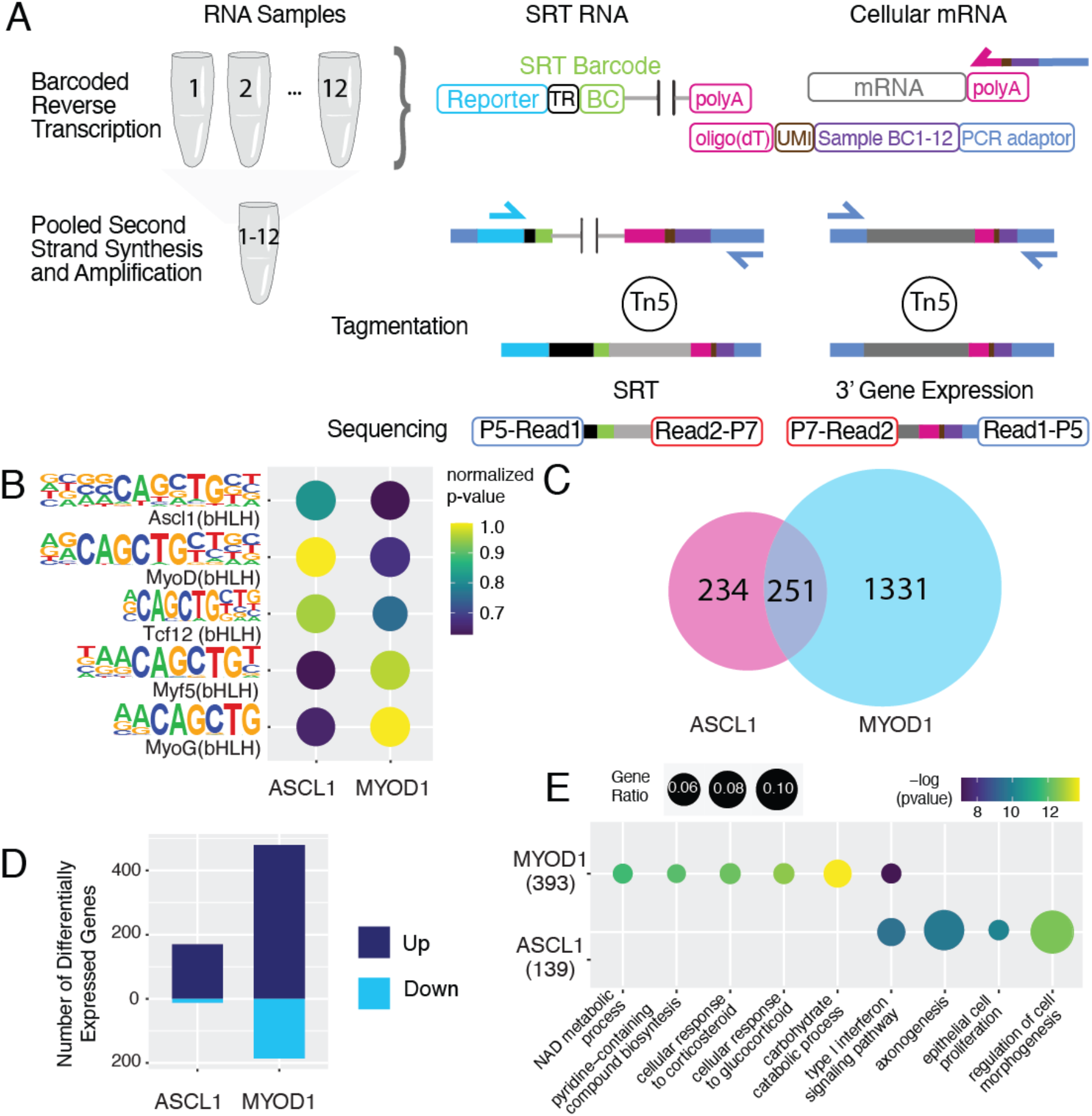
Barcoded SRT calling cards and transcriptomes enables joint measurement of TFBS and gene expression. A) Schematic overview of barcoded sequencing library preparation. Sample-specific barcode (Sample BC) with unique molecular identifiers (UMI) is introduced during reverse-transcription of poly(A) RNA including SRTs and mRNA. Reverse transcription products (cDNA) can then be pooled for second strand synthesis and amplification. Sequencing libraries are prepared for SRTs and transcriptomes in parallel. B) Barcoded SRT experiments recover binding motifs for ASCL1 and MYOD1. C) Venn Diagram showing shared and distinct genes near ASCL1 and MYOD1 binding sites. D) Transcriptomes profiled by bulk RNA-seq with barcodes revealed differential gene expression for ASCL1 and MYOD1, compared to cells transfected with unfused *piggyBac.* E) Gene Ontology of differentially expressed genes in ASCL1 and MYOD1 cells.

### Simultaneous Mapping of TFBS and Gene Expression of Pioneer TFs

ASCL1 and MYOD1 belong to a special class of transcription factors called pioneer factors that can access both open and closed chromatin and reprogram cell fate from pluripotent stem cells or fibroblasts to neurons and muscle cells respectively^19,35–37^. Chromatin immunoprecipitation followed by sequencing (ChIP-seq) after overexpression of these factors in mouse embryonic fibroblasts revealed a surprising degree of overlapping binding sites between these factors^18^. Gene expression profiling of TF overexpression, however, revealed differing transcriptional outcomes for these two factors^18^. As many TF binding events have no or small effects on gene regulation^38–41^, integrating TFBS data with mRNA-seq is a powerful method to decipher cis-regulatory modules and identify functional TFBS^40,42–44^. Typically, multi-omic measurements are collected from different populations of cells using separate protocols. In contrast, barcoded SRT calling cards and transcriptomes can be collected simultaneously from the same cells which may improve the ability to link TF binding to changes in gene expression.

As a proof-of-principle of this method, we transiently overexpressed unfused hyperactive *piggyBac* or fusions with ASCL1 or MYOD1 in HEK293T cells, then collected RNA after one week. We then prepared SRT calling card and poly(A) 3’ RNA-seq libraries in parallel. Cells transfected with ASCL1 and MYOD1 were co-transfected with non-overlapping pools of 12 barcoded SRTs to enable pooled SRT and transcriptome library preparation. Such experimental design therefore enables multiplexed TF profiling in a single pooled experiment. We performed 4 independent transfections for each factor. Compared to the recommended protocol for the original bulk RNA calling cards method for the same experiment^4^, barcoded SRT calling cards and transcriptomes reduces material cost and labor of experiments by over 10-fold (Table 1).

**Table 1:** Drastic cost and labor reduction of barcoded SRT and transcriptomes compared to original protocol. ‘Original’ calculations use the recommended 12 replicates per TF^4^. This experiment assayed 3 TFs (unfused hyper *piggyBac*, ASCL1, and MYOD1). Transfection costs are based on NEON or nucleofector transfection device reactions. Tagmentation costs assume a library is prepared for each of the 12 replicates for both calling cards and transcriptomes. Tapestation costs reflect core facility pricing.

| | Replicates (n) | | Cost (\$USD) | |
| --- | --- | --- | --- | --- |
|  | Original | Barcoded | Original | Barcoded |
| Transfections | 36 | 12 | 720 | 240 |
| RNA isolation and reverse transcription | 36 | 12 | 180 | 60 |
| Amplification | 72 | 2 | 216 | 6 |
| Bead Cleanup, Tapestation | 72 | 2 | 1080 | 30 |
| Tagmentation | 72 | 2 | 2160 | 60 |
| Bead Cleanup, Tapestation | 72 | 2 | 1080 | 30 |
| Sequencing | Same |  |  |  |
|  |  |  | 5436 | 426 |

Using the pooled barcoded SRT approach, we recovered hundreds of thousands of genomic insertion sites for each factor (Supplementary Table 3). Compared to unfused *piggyBac* binding sites, barcoded calling card peaks for ASCL1 and MYOD1 were again enriched for bHLH motifs including Ascl1 and MyoD (Fig. 4B). Comparing genes near identified TFBS, we again observed ASCL1 and MYOD1 had shared and distinct binding profiles (Fig. 4C) consistent with previous studies^17,18^. The genomic insertion sites recovered strongly overlapped those from our earlier experiments with unpooled sequencing library preparations, but the total number of sites was lower. This could reflect reduced library complexity after pooling. Future experiments to understand the decreased peak recovery would further improve this methodology.

Next, to identify transcriptional consequences of TF overexpression, we analyzed the gene expression profiles that were simultaneously captured with SRTs. Supporting the approach of pooling of barcoded first strands for 3’ gene expression library preparation, all 12 samples were well-represented in the sequencing data. Neither the average number of genes detected, nor the total RNA counts differed across factors and samples clustered by experimental condition (Supplementary Figure 3). We performed differential gene expression analysis on transcriptomes of cells transfected with ASCL1 or MYOD1 fusions compared against unfused *piggyBac* and identified 182 and 666 genes differentially expressed respectively. Of the differentially expressed genes, 170 and 480 were upregulated in ASCL1 and MYOD1 transfected cells respectively (Fig. 4D), consistent with known roles of these transcription factors as activators of gene expression. Gene Ontology analysis of upregulated genes recapitulated some relevant pathways in ASCL1 transfected cells, but many pathways were not related to neurogenic or myogenic pathways (Fig. 4E). Further, while some differentially expressed genes overlapped with genes near TFBS identified by barcoded SRTs, they were not enriched for such overlap. This is consistent with previous studies showing poor correlation between TF binding and gene expression^38–41^. Nevertheless, these results demonstrate a novel method to simultaneously collect TFBS and gene expression changes from the same SRT calling card experiment which may facilitate the inference of functional TFBS.

### Barcoded SRTs Facilitate in vivo Calling Card Experiments in Mouse Brain

We have previously demonstrated that SRT calling cards can be used to record TF binding *in vivo*^3,16^, though often requiring ~10 technical replicates per biological replicate. To test whether barcoded SRTs can also function *in vivo* and reduce this need for technical replicates, we performed calling card experiments with barcoded and non-barcoded SRTs in the mouse cortex. We packaged tdTomato SRT plasmids with or without barcodes as AAV and delivered them to cortex of mice as described^3,16^. Unfused *piggyBac* has an insertion preference at super-enhancers which are a class of enhancers regulating genes linked to cell identity^45,46^. Leveraging this property, calling cards have been used to read out these important regulatory elements^3,16,47^. To record these sites *in vivo*, we co-transduced mouse cortexes with unfused *piggyBac* and barcoded or non-barcoded SRTs.

After 21 days, we collected similar amounts of brain tissue from mice injected with barcoded or non-barcoded SRTs and prepared calling card libraries (Figure 5A). As with our *in vitro* experiments, all 25 unique barcodes were integrated into the genome and efficiently recovered (Fig. 5B). Lower recovery of 2/25 barcodes may reflect imbalances in vector DNA pooling prior to AAV packaging. After normalizing by the total depth of sequencing, we found that use of barcodes improved the recovery of SRTs and yielded around 2-fold more genomic insertions than non-barcoded counterparts (Fig. 5C). Genome-wide, integrations of barcoded and non-barcoded SRTs were highly concordant. Visualizing insertions and called peaks across the genome demonstrates this concordance (Fig. 5D). Analysis of genomic features of SRT insertion sites revealed similar insertional preferences (Fig. 5E). We recovered more insertions in promoter regions using barcoded SRTs, suggesting the unbarcoded SRTs might have had especially limited dynamic range in these regions. This would be expected as some of these loci are expected to contain strong binding sites or few ‘TTAA’ sequences, limiting the quantification of non-barcoded insertions.

**Figure 5:**
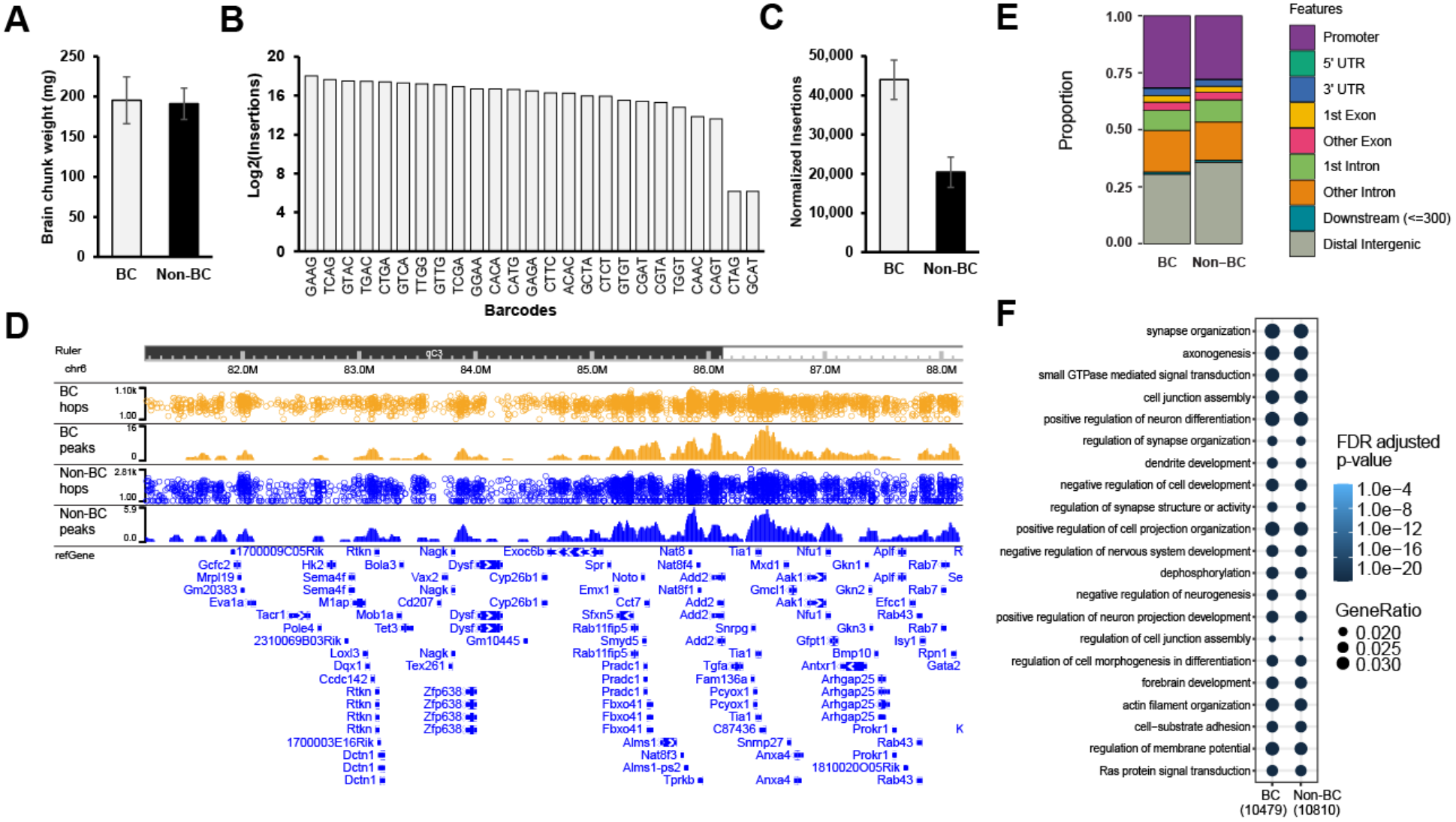
Comparison of barcoded and non-barcoded SRT calling cards *in vivo* in the mouse brain. A) Equivalent amounts of brain tissue were collected after *in vivo* calling card experiments using a pool of 25 barcoded (BC) or non-barcoded (non-BC) SRT donors delivered by AAV. n = 4 for BC and 3 for Non-BC. B) Number of genomic insertions recovered for each barcoded SRT. C) Number of genomic insertions recovered at the same depth of sequencing for barcoded and non-barcoded SRTs. D) Browser view of genomic insertions and called peaks for barcoded and non-barcoded SRTs. E) Genomic features of peaks called by barcoded and non-barcoded experiments. F) KEGG pathway enrichment comparison of genes near peaks called by barcoded and non-barcoded experiments.

Next, we performed functional enrichment analysis of genes located near insertions. Based on the tropism of AAV9, we expected the vast majority of insertions to be in neuronal cells, with some insertions in astrocytes^3,16^. Accordingly, barcoded and non-barcoded insertion sites were located near genes strongly enriched for neurological functions including synapse organization, forebrain development, and axonogenesis (Fig. 5F). Functional enrichment was similar for insertions of barcoded and non-barcoded SRTs and consistent with our previous findings^3,16^. Altogether, these results demonstrate that barcoded SRTs can recover biologically relevant binding events *in vivo* and outperform non-barcoded SRTs in the number of unique insertions at a fixed sequencing depth, while significantly reducing labor and reagent costs.

## Discussion

Understanding where TFs bind in the genome and how they orchestrate gene expression is a central goal in genomics^40,42–44^. Calling cards is a powerful functional genomics method to identify the binding sites of TFs and other chromatin-associated factors in mammalian cells both *in vitro* and *in vivo*^1–3,16,25^. The recently invented ‘self-reporting transposon’ converts the calling card recordings of TF binding to an RNA readout, enabling simultaneous profiling of gene expression and binding in single cells^3^. Here, we present two crucial modifications of the SRT technology and associated protocols to enable parallel recording of TFBS and gene expression in bulk populations of cells: barcoded SRTs and barcoded transcriptomes. Besides enabling transcriptomic measurement, these improvements also drastically reduce the experimental cost and labor of calling card experiments.

First, we performed targeted mutagenesis of the *piggyBac* transposon TR region. We coupled a simple PCR mutagenesis method with SRT calling cards to rapidly screen for positions in the *piggyBac* TR that could be mutated while retaining compatibility with transposition. Through this, we discovered four consecutive nucleotides within the TR that were tolerant of a range of mutations, both singly and in combination, without markedly reducing transposition efficiency. To our knowledge, these are the first reported mutations within the *piggyBac* terminal invert repeat that are compatible with transposition. We note the wild-type TR sequences are inserted at the highest frequency in our *in vitro* experiments compared to any 4-nt barcoded SRT so our targeted mutagenesis approach did not improve overall transposition efficiency. Nevertheless, for applications where the number of integrations is not paramount such as mutagenesis screening^48^, cellular lineage tracing^49^, and delivery of human gene therapy^50^, we anticipate that barcoded *piggyBac* transposons will have broad utility beyond calling card assays.

As a resource to the community, we have individually cloned the top 24 integration-competent error-detecting barcodes into two versatile self-reporting transposon vectors compatible with *in vitro* and *in vivo* experiments. Barcoded error-detecting SRT vectors will allow experimental conditions (e.g., timepoint, drug treatment) to be uniquely barcoded, pooled, and accurately demultiplexed without sample mis-assignment due to sequencing errors. Of note, multiple transcription factors can be assayed simultaneously in a pooled experiment by using non-overlapping sets of barcoded SRT for each TF.

We demonstrated that barcoded SRT calling cards identified TFBS for four bHLH transcription factors involved in cell fate specification and transdifferentiation: ASCL1, MYOD1, NEUROD2, and NGN1. We identified shared and unique binding sites for each factor and recovered binding motifs that matched known motifs for these factors. Supporting the identification of bona fide TFBS by barcoded SRT calling cards, we found that genes near TFBS were enriched for functions related to known functions of the assayed TFs. Barcoded SRT vectors reduced the experimental cost and labor of the calling card protocol by an order of magnitude which allowed us to easily measure TFBS for these four transcription factors in HEK293T cells. Remarkably, although HEK293T cells do not normally express any of the assayed TFs, all 4 TFs were able to recognize their cognate consensus motifs, and these motifs were located near genes with functions associated with each TF. The successful recovery of TFBS for all 4 TFs by barcoded calling cards demonstrates its versatility and its promise as an alternative to ChIP-seq. We also demonstrate that barcoded SRTs improve the recovery of integration events in the mouse cortex. This will facilitate future studies using calling cards to record TFBS *in vivo*. Combined with Cre driver lines, this method is also compatible with cell-type specific recording of TFBS in a mixed populations of cells *in vitro* or *in vivo*^16^.

Identifying TFBS is a first step toward understanding gene regulatory networks but many TF binding events have no or small effects on gene regulation^38–41^. Integrating multi-omics datasets, such as TFBS and mRNA-seq, is therefore necessary to identify functional TFBS governing gene expression^40,42–44^. Often, these multi-omics datasets are generated from different populations of cells using vastly different protocols that can introduce biases and batch effects. Since SRTs are expressed and collected as RNA, TFBS and gene expression data can be simultaneously generated from the same RNA sample using calling cards. While this method has been demonstrated in single-cell experiments^3^, bulk calling cards required modification to allow such joint measurement in bulk experiments. Specifically, we directly barcoded the SRT and introduced an additional barcode during reverse-transcription for barcoding mRNA. Combining barcoded SRT calling cards with bulk RNA barcoding and sequencing (BRB-seq) therefore enabled simultaneous identification of TFBS and gene expression in a protocol with drastically reduced cost and labor^13^.

We demonstrated that the combined protocol can jointly recover TFBS and gene expression during TF overexpression of the pioneer factors ASCL1 and MYOD1. Calling cards with barcoded SRTs and transcriptomes is therefore a novel and powerful method to infer functional TFBS in populations of cells. Technical and experimental optimizations of this method may improve its utility in future experiments. Further, using an inducible *piggyBac* system^24,51^ would enable temporal measurement of binding and expression changes. That application would be especially powerful during the time course of cellular reprogramming experiments to link TFBS to gene expression changes controlling cell fate specification.

Finally, our simple mutagenesis method will be useful for introducing barcodes to other DNA transposons such as *SleepingBeauty* and *Tol2*. Transposons are widely used for transgenics, mutagenesis, and functional genomics experiments^52^. As the SRT protocol can easily scale to recover millions of genomic integration sites, insertion preferences for other transposons can be readily ascertained using this method. Each transposon has its own preferences for genomic integration which can have complementary uses. Further, insertion profiles can depend on chromatin state^45^ so SRTs can potentially be used to read out chromatin status and histone modifications. For example, we have shown that calling card experiments using unfused *piggyBac* can identify super-enhancers^3,16,47^. Joint measurements of *piggyBac* insertions and gene expression with this method may help link super-enhancers to gene regulatory networks.

In conclusion, barcoded SRTs simplify bulk calling cards experiments, enable barcoding of experimental conditions, and allow for pooled library preparations that drastically reduce cost and labor. Incorporating barcoded transcriptomes into the library preparation enables joint measurement of transcription factor binding and gene expression from the same biological sample. This method will facilitate the inference of gene regulatory networks for TFs involved in development, cellular reprogramming, and disease.

## Acknowledgments

We thank Nancy Craig for useful discussions regarding *piggyBac* mutagenesis. We thank Jessica Hoisington-Lopez and MariaLynn Crosby from the DNA Sequencing Innovation Lab at The Edison Family Center for Genome Sciences and Systems Biology for their sequencing expertise. We thank Mingjie Li at the Hope Center Viral Vectors Core for AAV packaging services.

This work was supported by T32HL125241 (National Heart, Lung, and Blood Institute), T32HG000045 (National Human Genome Research Institute) (A.Y.), U54HD087011 (National Institute of Child Health and Human Development), R21NS087230-01A1 (National Institute of Neurological Disorders and Stroke) (R.D.M. and J.M.), RF1MH117070, RF1MH126723 (National Institute of Mental Health) (R.D.M., J.D.D.), R01GM123203 (National Institute of General Medical Sciences)(R.D.M.), SFARI Explorer 500661 (Simons Foundation Autism Research Initiative) (R.D.M., J.D.D.), R21 HG009750 (National Human Genome Research Institute) (R.D.M), P50 HD103525 (Eunice Kennedy Shriver National Institute of Child Health & Human Development) (J.M), and F30 HG009986 (A.M.). This work was further supported by the Hope Center Viral Vectors Core and a P30 Neuroscience Blueprint Interdisciplinary Center Core award to Washington University (P30 NS057105). GTAC@MGI is partially supported by National Institutes of Health (NIH) grants P30 CA91842 and UL1 TR000448. MAL is supported by the Seaver Foundation as a Seaver Faculty Scholar.

## Methods

### Transposon Mutagenesis

PCR mutagenesis was performed in a 50 μL reaction containing: 25 μl 2X Kapa HiFi HotStart ReadyMix, 1 μl of 10 μM SRT Mutagenesis Forward Primer (either puro or tdTomato version), 1 μl of 10 μM SRT Mutagenesis Reverse Primer, 100 ng of SRT DNA (either PB-SRT-puro or PB-SRT-tdTomato), and 22 μl of ddH2O. PCR reactions were performed following thermocycling parameters: 95°C for 3 minutes, 10 cycles of: 98°C for 20 seconds, 60°C for 30 seconds, 72°C for 2 minutes, then 72°C for 10 minutes, and 4°C forever.

PCR reactions were performed in duplicate. Each pool of mutant amplicons was purified with NucleoSpin Gel and PCR Clean-up (Macherey and Nagel). Products were transfected into separate wells of HEK293T cells to minimize any artifacts.

>SRT Mutagenesis Reverse Primer tgcatctcaggagctcttaacc**NNNN**aaagatagtctgcgtaaaattgac
>SRT Mutagenesis Forward (puro^R^) GCGGAAGGCCGTCAAGGCC
>SRT Mutagenesis Forward (tdTomato) CACGAGACTAGCCTCGAtcaaggcgcatttaaccctagaaagataa

### Cell Culture

HEK293T cells were maintained in Dulbecco’s Modified Eagle Media (DMEM) supplemented with 10% fetal bovine serum and 1% penicillin-streptomycin. Cells were passaged every 3–4 d by enzymatic dissociation using trypsin.

### Cloning

ASCL1, MYOD1, NEUROD2, and NGN1 were amplified from lentiviral cDNA expression vectors using 2X Kapa HiFi HotStart ReadyMix. A nuclear localization sequence was added to the 5’ end of each gene, and an L3 linker (amino acid sequence KLGGGAPAVGGGPKAADK) 24 was inserted between the TF and hyper-active *piggyBac.* EF1a_ASCL1, MYOD1, and NEUROG1_P2A_Hygro_Barcode were gifts from Prashant Mali (Addgene plasmid #120427, #120464, and #120467). phND2-N174 was a gift from Jerry Crabtree (Addgene plasmid #31822).

### Animals

All animal practices and procedures were approved by the Washington University in St. Louis Institutional Animal Care and Use Committee.

### Calling Card Experiments

Calling card experiments are performed as described with minor modifications^3,16,53^. Twenty-four hours before transfection, 250,000 HEK293T cells are plated per well in a 12 well plate. The next day, cells are transfected using PEI (Polysciences) with 1 ug of total DNA comprising 500 ng of *piggyBac* (fused or unfused) and 500 ng of donor SRT (purified PCR product or miniprepped DNA). Medium is changed 24 hours after transfection. Three days after transfection, each well is trypsinized and replated into a T25 flask. For puromycin-resistance SRTs, puromycin is added 24 hours later (2 ug / mL). Three days after puromycin selection, total RNA is harvested using Direct-zol RNA MiniPrep kit (Zymo Research). RNA concentration and integrity was assessed using Nanodrop.

### Virus Generation and Injections

Transposase and donor transposon constructs (barcoded or wildtype) were cloned into independent AAV transfer vectors and used for *in vitro* transfection or viral packaging. Plasmids were packaged into AAV9 by the Hope Center Viral Vectors Core at Washington University School of Medicine. For *in vivo* experiments, intracranial injections into the cortex of wildtype C57BL6/J P0-1 mice of both sexes were performed as previously described^16^. 1ul of viral mix was delivered across 3 sites per hemisphere for a total of 6ul per brain. Viral titers (viral genomes [vg] per milliliter) were ~1.0 × 10^13^. Animals were euthanized at P21 for analysis, cortices were dissected, and RNA extraction was conducted as described^16^.

### Calling Card Library Preparation

Calling card libraries were prepared as described with minor modifications^3,4,16^. We performed first-strand reverse transcription reactions in 20 μL total volume using 2 μg of RNA from each *in vitro* sample and 4 ug from each *in vivo* sample. RNA mixed with water and dNTPs was hybridized to oligo-dT primers (1 μL of 50 μM SMART_dTVN) by incubation at 65 °C for 5 minutes and immediately transferred onto ice. 0.5 μL of Maxima H Minus Reverse Transcriptase, RNasin RNAse inhibitor, and 5X RT buffer were added and samples are incubated at 50 °C for 60 min for reverse transcription.

### Barcoded Calling Card and Transcriptome Library Preparation

Bulk RNA Barcoding and sequencing (BRB-seq) was performed with minor modifications^13,32^. We performed first-strand reverse transcription reactions in 20 μL total volume using 2 μg of RNA from each sample. Barcoded BRB-seq_dT30VN primers modified to mimic the 10x Genomics v2 chemistry. RNA mixed with water and dNTPs was hybridized to barcoded oligo-dT primers (2 μL of 25 μM stock) by incubation at 65 °C for 5 minutes and immediately transferred onto ice. 1 μL of template switch oligo (TSO_SMART), 0.5 μL of Maxima H Minus Reverse Transcriptase, RNasin RNAse inhibitor, and 5X RT buffer were added and samples were incubated at 50 °C for 60 min for reverse transcription.

For barcoded SRT and transcriptome experiments studying ASCL1, MYOD1, and unfused *piggyBac*, 3 μL of reverse-transcription product from 4 replicates for each factor were pooled together for transcriptome analysis. 4 replicates of each factor (5 μL per replicate) were pooled and purified in parallel for calling card library preparation. Pooled samples were purified using NucleoSpin Gel and PCR Clean-up (Macherey and Nagel) and eluted with 30 μL of elution buffer. We designed barcoded oligoDT-VN oligos to mimic 10x Genomics v2 chemistry (partial seq1, 16 bp cell barcode extracted randomly from the 10x Genomics safelist, and a 10 bp UMI (5N + 5V). Barcoded primer sequences are provided in Supplemental Table S1.

Barcoded, pooled first-strand reactions (23 μL) were mixed with 1 μL partial seq1 primer (10 μM), 1 μL SMART primer (10 μM), and 25 μL 2X KAPA HiFi HotStart ReadyMix (Roche). 10 cycles of PCR with a long extension time (98° 20 seconds, 60° 30 seconds, 72° 6 minutes) were performed.

cDNA was purified with 0.6X AMPure XP (Beckman Coulter) magnetic beads. DNA was eluted with 20 μL water and concentration was measured using the Tapestation D5000 ScreenTape (Agilent). 600 pg of product were tagmented with barcoded N7 primers and P5-index-seq1 primers using the Nextera XT kit (Illumina). BRB-seq libraries were sequenced on a Novaseq 6000 paired-end with 28 x 91 reads.

### Primers

>SMART_dT18VN AAGCAGTGGTATCAACGCAGAGTACGTTTTTTTTTTTTTTTTTTTTTTTTTTTTTTTVN
>BRB-seq_dT30VN CTACACGACGCTCTTCCGATCT<u>CTGATAGCATGGTCAT</u>NNNNNVVVVVTTTTTTTTTTTTTTTTTTTTTTTTTTTTTTVN
>SRT_PAC_F1 CAACCTCCCCTTCTACGAGC
>SRT_tdTomato_F1 TCCTGTACGGCATGGACGAG
>SMART_TSO AAGCAGTGGTATCAACGCAGAGTACrGrGrG
>SMART AAGCAGTGGTATCAACGCAGAGT
>Partial Seq1 CTACACGACGCTCTTCCGATCT

Barcoded *piggyBac* primers, for example:

>OM-PB-ACG (barcode sequence is underlined) AATGATACGGCGACCACCGAGATCT[optional_index]ACACTCTTTCCCTACACGACGC TCTTCCGATCT<u>ACG</u>CGTCAATTTTACGCAGACTATCTTT
>P5-index1-Seq1 (index sequence is underlined) AATGATACGGCGACCACCGAGATCTACAC<u>AGGACA</u>ACACTCTTTCCCTACACGACG CTCTTCCGATCT
>Nextera_N70X (index sequence is underlined) CAAGCAGAAGACGGCATACGAGAT<u>TCGCCTTA</u>GTCTCGTGGGCTCGG

### Library Prep and Sequencing

Purified PCR product was measured using a D5000 ScreenTape (Agilent). cDNA samples were diluted to 600 pg/μL and 2 μL of this was used for tagmentation with Nextera XT kit.

Agencourt Ampure XP (Beckman Coulter)

Nextera XT DNA Library Preparation Kit (Illumina, Inc)

High Sensitivity D1000 ScreenTape (Agilent Technologies)

### RNA-seq Analysis

Sequencing data corresponding to barcoded bulk RNA transcriptomes were processed using the 10x Genomics software package Cell Ranger (v 2.1.0). The output filtered gene expression matrices were imported into R (v 3.5.1) for further analysis^54^. Gene counts were used directly in edgeR for standard bulk RNA-seq analysis^55^.

### Calling Card Analysis

#### Sequencing and analysis

Bulk barcoded RNA calling card libraries were sequenced and analyzed as described with modifications to utilize the SRT barcode^3^. Calling card reads begin with a 3-nucleotide library barcode, the barcoded transposon TR, the insertion motif TTAA, then the genome at the site of insertion. Reads are checked for the library barcode, TR sequence, and TTAA and these sequences are trimmed. SRT barcodes are extracted by UMI-tools^56^, and appended as a sequence tag to the read. Any remaining Nextera adaptors are trimmed before mapping the reads to the human genome (hg38) using NovoAlign. Aligned reads are validated as insertions if adjacent to a TTAA site in the genome. Bona fide insertions are then converted to qBED format (née .ccf)^57^. SRT barcodes were incorporated into the barcode column of the qBED. If non-overlapping barcode sets were used to define experiments, qBED files can be demultiplexed by this barcode field.

Raw and processed sequencing data generated in this study has been submitted to the NCBI Gene Expression Omnibus (GEO; https://www.ncbi.nlm.nih.gov/geo/) under accession number GSE195992. All software used to perform the analyses are available at:

SRT Calling card tools: https://github.com/arnavm/calling_cards
Barcoded SRT Calling cards tools: https://github.com/aMattScientist/barcoded_calling_cards
Peak Calling CCF tools: https://gitlab.com/rob.mitra/mammalian_cc_tools

#### Peak calling

Calling card peaks were called as described ^1,25^ using in-house peak calling software. Specifically, peaks were called using the call_peaks_macs python script, which follows the algorithm used by MACS to call ChIP-Seq peaks^58^ modified for the analysis of calling card data. The main peak calling function is passed an experiment frame, a background frame, and an TTAA_frame, all in qBED/ccf format^57^. It then builds interval trees containing all of the background and experiment hops (insertion events) and all of the TTAAs. Next, it scans the genome with a window of window_size and step size of step_size and looks for regions that have significantly more experimental hops than background hops (poisson w/ pvalue_cutoff). It merges consecutively enriched windows and computes the center of the peak. Next it computes lambda, the number of insertions per TTAA expected from the background distribution by taking the max of lambda_bg, lamda_1, lamda_5, lambda_10. It then computes a *p*-value based on the expected number of hops = lamda * number of TTAAs in peak * number of hops in peak. Finally, it returns a frame that has Chr, Start, End, Center, Experiment Hops, Fraction Experiment, Background Hops, Fraction Background, Poisson *p*-value as columns. We used parameters: -pc 0.001 --peak_finder_pvalue 0.01 --window 1000 --step 500 --pseudocounts 0.2 for peak calling.

