## Supplementary Material for "Measuring Transcription Factor Binding and Gene Expression using Barcoded Self-Reporting Transposon Calling Cards and Transcriptomes"

### Supplementary Material for Lalli *et al.* Barcoded Calling Cards

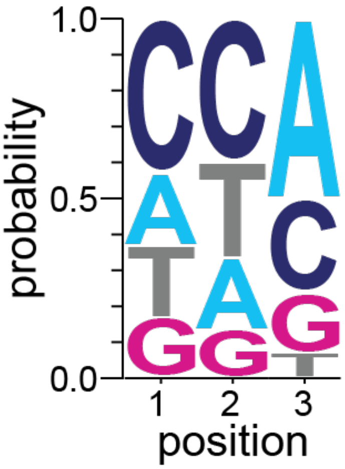


**Supplementary Figure 1:** Sequence logo of the top 30 most abundantly inserted 3-nt barcoded SRTs reveals modest sequence preference for integration efficiency.

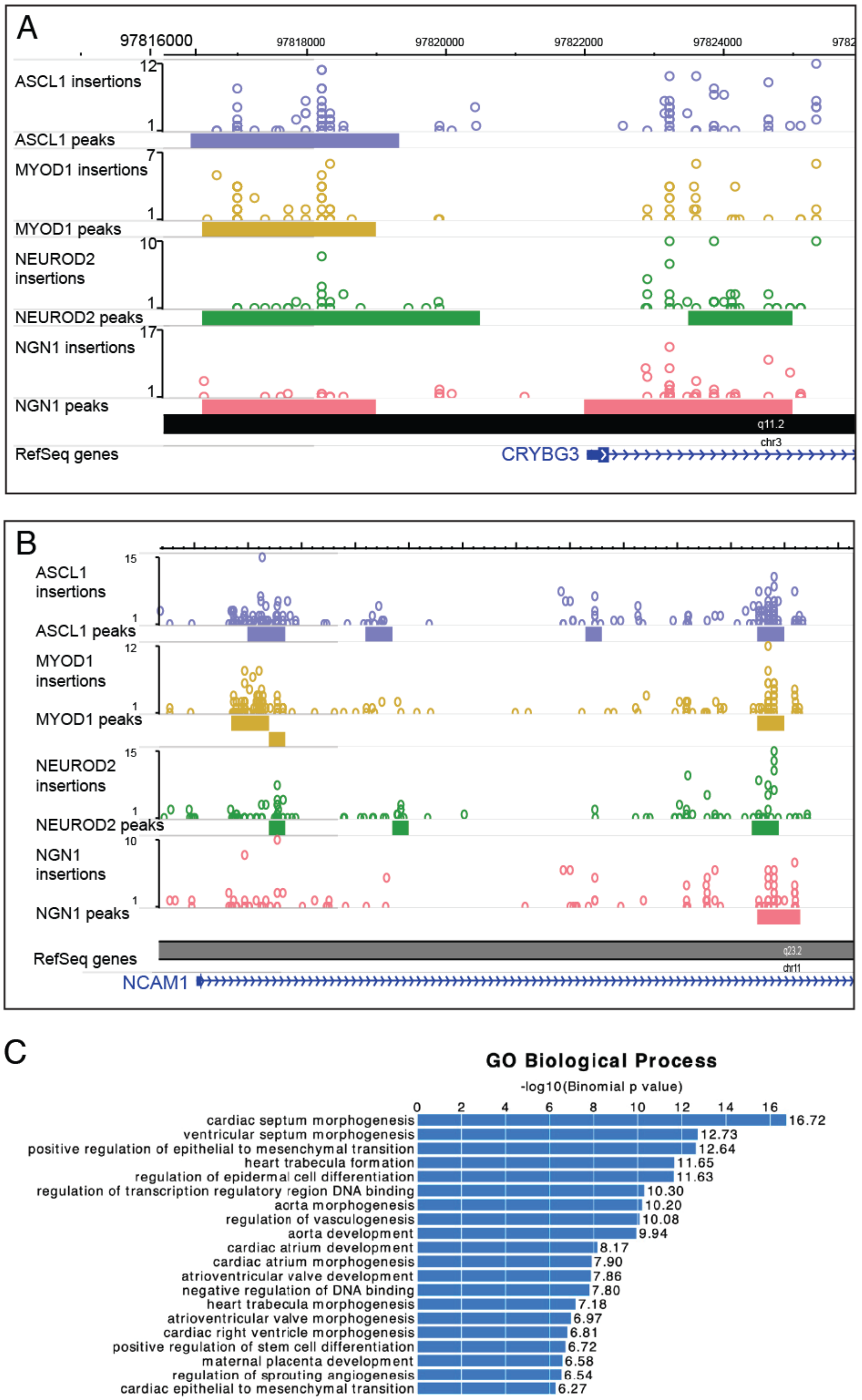


**Supplementary Figure 2:** Calling card insertions and peaks for four transcription factors at shared and distinct genes. Browser views of genomic insertions and called peaks highlight binding sites shared across all factors at A) *CRYBG3* and B) *NCAM1*. C) GREAT analysis of MYOD1 hops identifies pathways enriched in heart development.

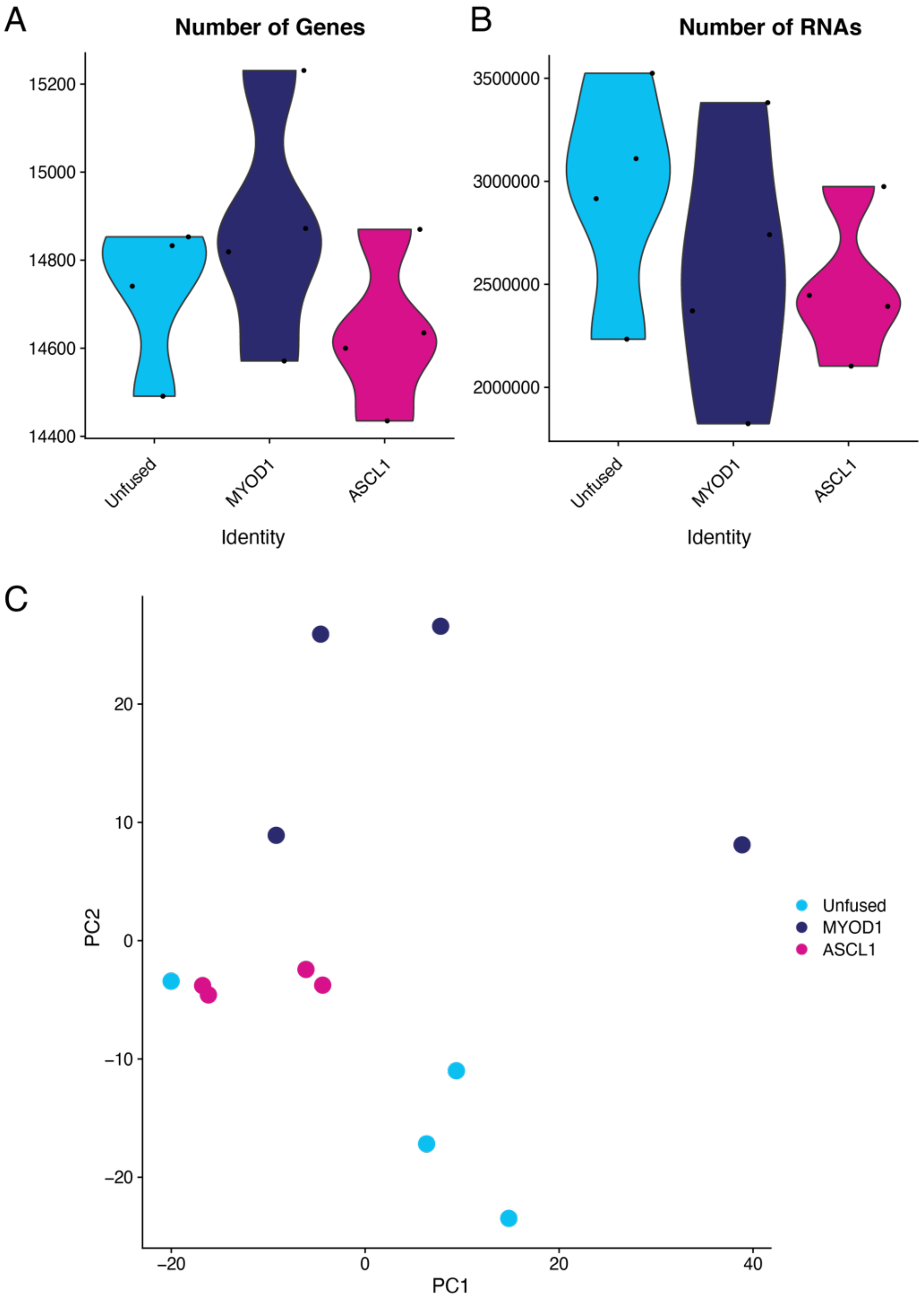


**Supplementary Figure 3:** Quality control of Bulk Barcoded RNA sequencing samples. A) Number of genes detected in each sample. B) Total number of RNAs collapsed by unique molecular identifiers. All 12 samples, which were prepared and sequenced as a single pool, were present and contained approximately the same number of genes and RNAs detected. C) Principal component analysis for dimensionality reduction and visualization of samples shows clustering by experimental subgroup.

| *Sequence* | *Number of Events* |
|---|---|
| *TT*CTAGGG* | 9256 |
| TT**C**CTAGGG | *6430* |
| *TT**C**CTAGG_* | 5476 |
| *TT*CTA**C**GG* | 3878 |
| *TT*CT**C**GGG* | 2870 |
| TT**G**CTAGGG | *2748* |
| *TT*CT**G**GGG* | 2570 |
| *TT*CTA**A**GG* | 2202 |
| *TT*CT**T**GGG* | 2019 |
| *TT*CTA**T**GG* | 1887 |
| TT**G**CTAGG_ | *1625* |

**Supplementary Table 1:** piggyBac TR does not accommodate a 1-nt insertion at position marked by asterisk (*). Wild-type sequence (red). Sequences with 1-nt insertion (italics) also had 1-nt deletion in 2/3 cases. Mutations to wild-type sequences are **bolded.** Underlines mark single nt deletions.

| Replicate | Transcription Factor | Self-Reporting Transposon | Hops | Total Hops | Called Peaks |
|---|---|---|---|---|---|
| 1 | Unfused hyperPBase | WT PB-SRT-Puro | 191776 | | |
| 2 | Unfused hyperPBase | WT PB-SRT-Puro | 161056 | | |
| 3 | Unfused hyperPBase | WT PB-SRT-Puro | 222416 | 643632 | N/A |
| 1 | NEUROD2 | WT PB-SRT-Puro | 201986 | | |
| 2 | NEUROD2 | WT PB-SRT-Puro | 192798 | | |
| 3 | NEUROD2 | WT PB-SRT-Puro | 213119 | | |
| 4 | NEUROD2 | WT PB-SRT-Puro | 245030 | 852501 | 4594 |
| 1 | ASCL1 | Barcoded PB-SRT-tdTomato | 207857 | | |
| 2 | ASCL1 | Barcoded PB-SRT-tdTomato | 207771 | | |
| 3 | ASCL1 | Barcoded PB-SRT-tdTomato | 280185 | | |
| 4 | ASCL1 | Barcoded PB-SRT-tdTomato | 316377 | 1011703 | 4027 |
| 1 | MYOD1 | Barcoded PB-SRT-tdTomato | 305021 | | |
| 2 | MYOD1 | Barcoded PB-SRT-tdTomato | 302416 | | |
| 3 | MYOD1 | Barcoded PB-SRT-tdTomato | 246855 | 853965 | 6178 |
| 1 | NGN1 | Barcoded PB-SRT-puro pool | 189488 | | |
| 2 | NGN1 | Barcoded PB-SRT-puro pool | 216185 | | |
| 3 | NGN1 | Barcoded PB-SRT-puro pool | 254215 | | |
| 4 | NGN1 | Barcoded PB-SRT-puro pool | 249453 | 904991 | 3593 |

**Supplementary Table 2:** Description of Calling Card Experiments. All experiments had ≥ 3 replicate transfections. Self-reporting transposon reporter gene is indicated. Number of genomic insertions (hops). N/A, not applicable. Genomic insertions of unfused hyper-piggyBac are used as the background to call peaks. per replicate are indicated. Cumulative number of hops for each transcription factor, and number of called peaks are shown.

| Transcription Factor | Self-Reporting Transposon | Reads | Uniquely Mapped | Hops | Peaks |
|---|---|---|---|---|---|
| ASCL1 | Barcoded PB-SRT-puro pool | 3865901 | 3191357 | 460634 | 526 |
| MYOD1 | Barcoded PB-SRT-puro pool | 3083682 | 2823939 | 510573 | 1953 |
| Unfused | Barcoded PB-SRT-puro pool | 3513592 | 2823939 | 312024 | N/A |

**Supplementary Table 3:** Description of Calling Card and Transcriptome Experiments. Self-reporting transposon reporter, number of total sequencing reads, number of uniquely mapped reads, and number of genomic insertions (hops) are indicated. N/A, not applicable. Genomic insertions of unfused hyper-piggyBac are used as the background to call peaks.
